# Understanding metabolic behaviour in whole-cell model output

**DOI:** 10.1101/2020.08.19.257147

**Authors:** Sophie Landon, Oliver Chalkley, Gus Breese, Claire Grierson, Lucia Marucci

## Abstract

Whole-cell modelling is a newly expanding field that has many applications in lab experiment design and predictive drug testing. Although whole-cell model output contains a wealth of information, it is complex and high dimensional, thus hard to interpret. Here, we present an analysis pipeline that combines machine learning, dimensionality reduction and network analysis to interpret and visualise metabolic reaction fluxes from a set of single gene knockouts simulated in the *Mycoplasma genitalium* whole-cell model. We found that the reaction behaviours show trends that correlate with phenotypic classes of the simulation output, highlighting particular cellular subsystems that malfunction after gene knockouts. From a graphical representation of the metabolic network, we saw that there is a set of reactions that can be used as markers of a phenotypic class, showing their importance within the network. Our analysis pipeline can support the understanding of the complexity of *in silico* cells without detailed knowledge of the constituent parts, which can help to understand the effects of gene knockouts, and, as whole-cell models become more widely built and used, aid genome design.

## Introduction

Recent years have seen a significant increase in the availability of high-throughput biological data (Gomez-Cabrero et al. 2014). Methods for collecting cellular data such as genome sequences (Sanger sequencing (Sanger et al. 1977), DNA microarrays (Southern 2001) and next-generation sequencing (Schuster 2008)), RNA transcripts (microarray sequencing (Stark et al. 2019)) and protein concentrations (mass spectrometry (Aebersold & Mann 2016)) are becoming cheaper and more accessible (Wetterstrand 2010). The integration of these data types reveals interactions between cellular processes (Manzoni et al. 2018), aiding analysis (Zampieri et al. 2019); also, significant leaps in the scale and capabilities of biological modelling give great scope for *in silico* data generation. Although mathematical models can never fully replicate living cells, their output can help to understand biological mechanisms and inform experimental design to improve *in vivo* data collection. These models can formalise processes at a specific level (e.g. translation) or construct a trans-omic network of the relationship between different cellular processes (Yugi et al. 2019) and couple metabolism with gene expression (O’brien et al. 2013).

Whole-cell models are the largest scale, and simulate every cellular process throughout the life cycle of a cell — only two are published, which model the life cycle of *Mycoplasma genitalium* (Karr et al. 2012) and *Escherichia coli* (Macklin et al. 2020). We focus on the *M.genitalium* model, which consists of 28 submodels that use a range of mathematical methods (linear programming, differential equations, geometry) to represent processes such as metabolism, RNA decay and cytokinesis, which integrate together at every time step. It contains thousands of parameters found through an intensive literature search or chosen to be biologically realistic — some parameters come from different organisms, as data gathered from *M.genitalium* is less ubiquitous than from more widely studied bacteria. The model is highly complex, computationally expensive, and generates huge amounts of time series data relating to thousands of variables. Interpreting whole-cell model *in silico* data can be difficult, but large-scale analysis is possible.

To interpret these data, tools and software are required to automatically process and consolidate the output so they can be viewed and interpreted easily, even by those with little computational expertise. Existing software tools that visualise whole-cell model output (Karr & Pochiraju 2018) handle one simulation at a time or multiple time series (Lee et al. 2013), but their capacity for processing large and varied datasets is limited — they focus on visualisation of different output streams, so all analysis is done by eye and there is no dimensionality reduction or statistical methodology.

A whole-cell model, with appropriate analysis software to process its output, could be a powerful predictive tool for gene editing. Genetic modifications can be trialled in a model before being physically made, which may save time and resources, and whole-cell models can be coupled with algorithms to predict gene knockouts intended to produce a chosen phenotype (Haimovich et al. 2015). As whole-cell models generate large datasets, machine learning methods are suitable for their analysis. These methodologies are data-driven, so can identify correlations and classify data with few assumptions and little biological knowledge. They are suited to biological model output, as the mechanisms for modelling individual subsystems are well established, but the relationships between different subsystems are not always clear.

Constraint-based metabolic models (CBMs) calculate the flow of substrates through the network of chemical reactions making up a cell’s metabolism (Bordbar et al. 2014). Each model consists of a stoichiometric matrix, where columns correspond to reactions, and rows to substrates, with upper and lower bounds for the rate of each reaction. These models are widely used to predict steady state metabolic fluxes, including the phenotypes of gene knockouts and synthetic lethal pairs and triplets (pairs or triplets of genes that when knocked out together cause cell death, but when knocked out singly do not) (Pratapa et al. 2015). CBMs do not provide unique solutions to the constraint-based problem — the optimal flux vector is chosen based on the objective function, which can vary with different models. The removal of reactions through gene knockouts can also result in re-routing of flux through different pathways (Güell et al. 2014), showing behaviour that may be very different from a wild type cell, but nonetheless still results in growth and division.

Metabolic modelling has also been integrated with transcriptional processes (King et al. 2015), tRNA charging and translation, to create ME-models (metabolism and expression) which tend to either equal or improve on CBMs accuracies (Lloyd et al. 2018). They contain some functional gaps, such as regulatory mechanisms (O’brien et al. 2013), but they are important steps towards whole-cell models, which couple multiple sub-models to simulate a cell life cycle (Karr et al. 2012).

There have been many applications of machine learning to CBMs, consolidated by Zampieri *et al.* (Zampieri et al. 2019), which show a range of uses for machine learning with biological modelling. Many have coupled CBMs with discriminative classifiers such as support vector machines (algorithms that define a hyperplane to separate different classes (Noble 2006)), neural networks (a system of connected nodes that can be trained to classify things by recursively adjusting the weights of connection between the nodes (Yegnanarayana 2009)), or random forests (a collection of decision trees that classify through finding the mean prediction from all the trees (Ho 1995)), to predict or classify gene essentiality, drug side effects, and protein functions. Others have used unsupervised learning to explore patterns and pathways in metabolic systems. Additionally, Rana *et al.* proposed two workflows for the coupling of machine learning and CBMs — the first where machine learning can be used to reduce the complexity of CBMs by finding the most important features of the model that can still match experimental data, and the second where machine learning can be used to tune CBMs to match their output more closely to experimental data (Rana et al. 2020).

These methods of prediction and analysis can be scaled to whole-cell models. However, whole-cell model output is composed of time series — contrary to CBM and ME output, which are steady-state rates or concentrations — and the labelling of these types of data is increasingly becoming a barrier to large-scale machine learning. As computational power increases and new data analysis algorithms are developed, the availability of fully labelled datasets to train and validate models is a limiting factor, and so new methods are being formed to automatically label data.

Time series data comes from all physical systems. Difficulty in interpreting it arises from the importance of ordering of different events, meaning that attributes of the data are dependent on each other in complex ways (Hannan 2009). Examples of machine learning methods applied to time series classification include music structure analysis (Serra et al. 2014), driver detection (Bernardi et al. 2018) and epileptic brain activity (Chaovalitwongse et al. 2006), using support vector machines, dynamic time warping and neural networks. Of the various methods available, deep learning has emerged as the most reliable (Fawaz et al. 2019, Wang et al. 2017), although accuracies of each method vary with different datasets. There are also other factors that affect the performance of an algorithm, such as feature selection and data pre-processing.

Many of these methods are supervised, meaning that they require labelled data in order to train a model. Historically, these labels would be manually generated by an expert to capture the ground truth of the problem. However, labelling data manually is time consuming and unfeasible for huge datasets. A solution to this problem is weak supervision, which uses weak labels (that do not express the ground truth) created from a model designed to map labels onto instances of the data (Zhou 2018). Snorkel is a methodology that creates a generative model (a statistical model of the joint probability of a variable and target label) to automatically produce weak labels, after collating metrics from multiple manually defined labelling functions (Ratner et al. 2019, 2017).

Although machine learning is a broadly used tool for classification and prediction, most algorithms are treated as black boxes and so results are created without context. For explanations of the functions of underlying structure in complex systems, network science can be used (Gosak et al. 2018). Network science is an area that has long been applied to analysis of biological systems: protein interactions, metabolic reactions and transcription regulation can be formalised as networks, leading to discoveries regarding properties of their interactions (Barabasi & Oltvai 2004). Network structure has been used to predict metabolic functions and find pathways for metabolite flow (Stelling et al. 2002), and to find control loops within gene networks (Wong et al. 2012).

The complexity of genomic interactions, even in cells as small as *M.genitalium*, is such that there is not necessarily a clear path from the genome after knockouts to the end phenotype. Even with functional annotations of genes, the genomic context of the remaining genome (which will be several hundred genes after a single gene knockout) cannot be disregarded as there may be redundancy in the genome, or unprecedented gene product interactions. The removed gene/s will not tell the full story, but zooming out to examine a large set of different genotypes through their metabolic fluxes can show us the trends across the full set of knockouts, providing a different angle than that of focusing on a single gene.

Here, we present a novel analysis pipeline which combines whole-cell model simulations of wild-type and gene knockout cells with time series classification and network analysis. The main steps automatically label a large dataset of metabolic fluxes, classify the flux behaviour as normal or abnormal (with respect to wild-type simulations), visualise the data and analyse the network topology. A schematic of our pipeline is shown in Figure 1; the main steps include automatic labelling of the fluxes as normal or abnormal (where normality refers to the behaviour of a reaction flux from a knockout simulation with respect to the behaviour of that reaction in a wildtype simulation), dimensionality reduction of the reactions for visualisation, and network analysis of the reactions. This analysis — looking at intermediate steps that connect genotype to phenotype — aims to increase our understanding of cellular processes, as well as provide foundations for *in silico* genome design.

**Figure 1:**
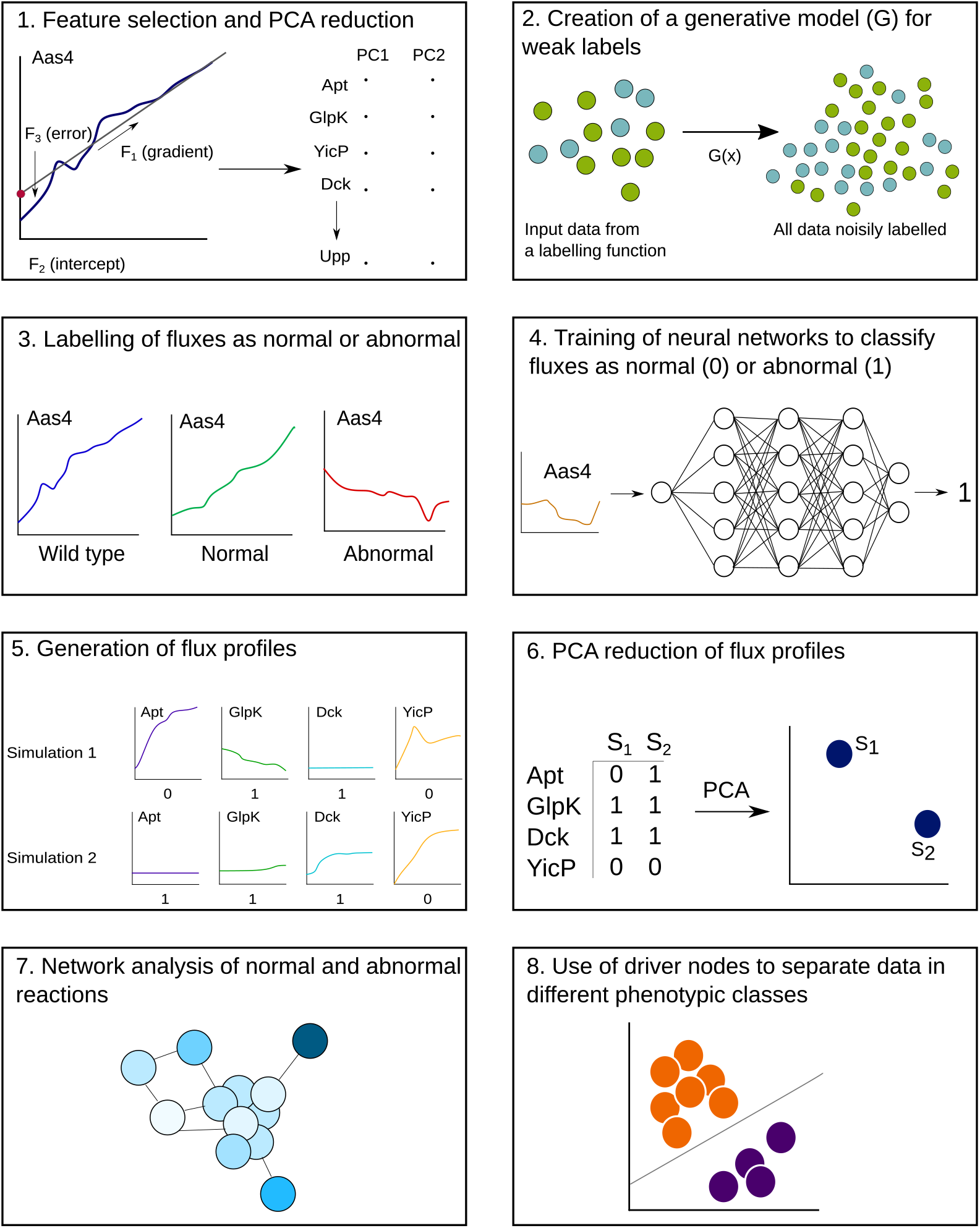
Step-by-step workflow of the analysis pipeline, beginning with the metabolic fluxes from the whole-cell model output. Steps 1-4 are applied to a training dataset of gene knockout simulations, where the end result is a trained neural network for each reaction. 1) For each reaction flux time series, four features are extracted and reduced to two dimensions through PCA. 2, 3) The extrema of these data are used to define boundaries for normal and abnormal behaviour, which are then used to create a generative function to map labels onto the reactions. 4) Neural networks are trained using this labelled data to classify reactions as normal or abnormal. Steps 5-8 are applied to a separate analysis dataset of gene knockout simulations. 5) The neural networks are used to classify the analysis dataset and create a flux profile for each simulation. 6) The flux profiles are reduced to two dimensions and plotted. 7, 8) Network analysis of the reactions reveals nodes that control the metabolic network, and correlate with different phenotypes after gene knockouts.

## Results

For large-scale analysis of the *M.genitalium* model output, we required two datasets: one for training the neural networks (box four in Figure 1) which required automatic label generation though Snorkel (boxes one and two in Figure 1), and one for analysis of the single gene knockouts (boxes 5-8 in Figure 1). The first (training) dataset contained three replicates each of all 359 possible single gene knockouts, plus 200 wild-type simulations, giving 1270 simulations in total. Each simulation had 279 dynamic reactions that vary in time (many reactions were consistently at steady state throughout the cell life cycle, which does not require a complex classifier to identify), with up to 50000 timesteps of a second. This was split into 80% training data, and 20% test data to validate the neural networks. The second (analysis) dataset consisted of ten replicates each of the 359 knockouts, with the same number of reactions and timesteps, totalling 3411 simulations. There are some gaps in the dataset, as some files were corrupted, and one knockout consistently caused the model to crash.

The *M.genitalium* whole-cell model has drastically varying fluxes through different reactions (see Figure S1). Furthermore, it is not always clear how the removal of a particular gene will affect cellular processes or cell viability. For each reaction, we presume there is a range of normal behaviour over which the cell can produce all necessary compounds for cell division, and dynamics outside of that range result in negative effects (e.g. build up or depletion of certain metabolites) that would affect the rest of the metabolism, and disrupt other processes, potentially causing cell death. The normality of reaction fluxes in a simulation can be used to understand the effects of gene knockouts through the cell cycle, and how metabolism is affected. This can help with predicting and explaining the effects of gene knockouts, and looking at patterns seen across different simulations. We visualised the reaction flux behaviour across our entire dataset, and looked at the topology of the metabolic network (in particular, how the network can be controlled by input nodes) to help explain the role of different reactions.

### Implementation of Snorkel for weak labelling

Manual labelling was not practical with such a large dataset, so we implemented Snorkel, which has been shown to perform as accurately as hand labelling in a previous study (Ratner et al. 2019). Snorkel requires manually defined labelling functions, which are an important heuristic for the basis of the generative model. The underlying patterns are used to form probabilistic labels, so together they need to capture some approximation of ground truth. In this case, we created labelling functions by amalgamating four key features extracted from each reaction flux time series. The trends in the different flux behaviours across the dataset were that the fluxes could be increasing, decreasing, or flat, and either relatively smooth or significantly oscillatory. A linear regression function was fitted to each time series, and the intercept, gradient, coefficient of determination, and mean squared error were found (Figure 2a). These captured the variation observed: smoothness in the mean squared error, the increasing or decreasing nature in the gradient, and the linearity in the coefficient of determination.

**Figure 2:**
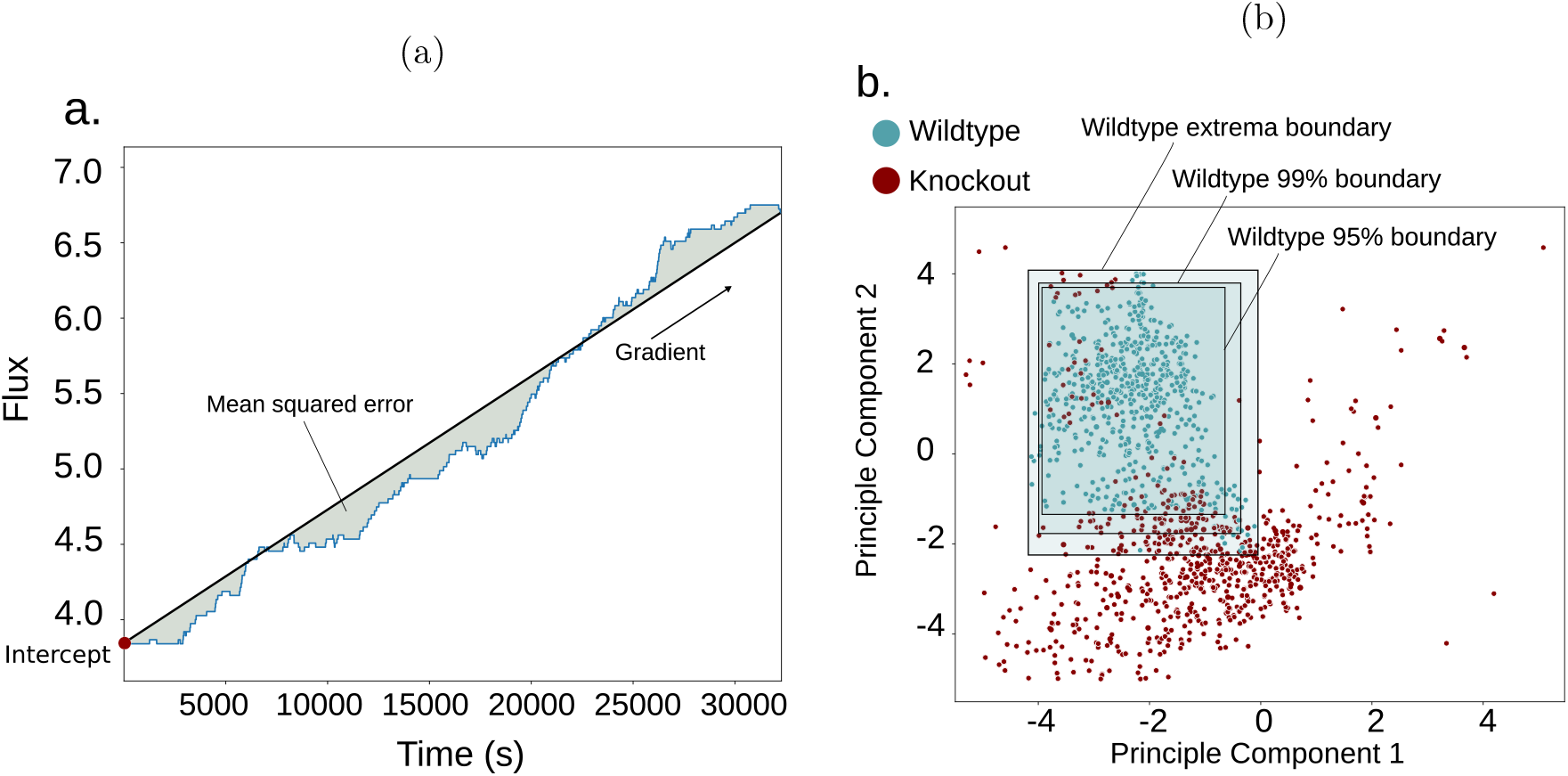
Method of feature extraction and normality classification shown graphically. The features shown in plot (a) were taken and PCA was applied for each reaction flux in each simulation across the dataset to create plot (b). The wildtype simulations are shown in blue, and then three boundaries are shown (extrema, 99% confidence intervals, and 95% confidence intervals), which are used to form three labelling schemes.

The results were reduced to two dimensions through principle component analysis (retaining an average of 82.9% of the variance across the reactions), leaving a 2D space over which boundaries of different thresholds could be drawn. Loosely, the boundaries were defined by the extrema of the wild-type simulations, which were taken to be the edges of normal behaviour for each reaction (Figure 2b). Any simulations outside these boundaries were classified as abnormal.

Three different boundaries were defined for different labelling schemes, as different confidence thresholds performed better or worse depending on the reaction. Boundaries at the extrema, then at 99% and at 95% were selected as the three labelling functions after comparison of their performance, and then combined to form the generative model. We then implemented Snorkel, leaving us with 1270 weakly labelled time series for each reaction. Ten reactions were manually labelled as normal or abnormal to test the accuracy of Snorkel’s labels, where characteristics like smoothness or the increasing or decreasing nature of the time series were used as comparison features to decide whether the behaviour of a reaction was normal or abnormal. The majority of the Snorkel labels gave over 90% accuracy, with the lowest at 69.2% (see Methods and Table 1).

**Table 1:** Accuracies of Snorkel’s weak labels for ten manually labelled reactions.

|  | Aas4 | AceE | Adk3 | Apts_Asp | Apts_Trp | ArcC | DcdK | Pyk_DADP | Pyl_GDP | TX_AROP22 |
| --- | --- | --- | --- | --- | --- | --- | --- | --- | --- | --- |
| Accuracy | 99.6% | 99.1% | 90.2% | 95.8% | 83.1% | 77.8% | 97.0% | 85.0% | 69.2% | 94.3% |

### Training of neural networks and flux profiling

The Snorkel results were used to train a neural network for each reaction. Although many machine-learning discriminative algorithms can be used for classification, artificial neural networks are some of the most effective (Caruana & Niculescu-Mizil 2006, Raczko & Zagajewski 2017). Neural networks consist of layers of nodes, representing artificial neurons with assigned weighted connections. The weights are adjusted through rounds of backpropagation or epochs until they predict correct classes for different types of input (Kröse et al. 1993).

Once trained (see Methods section), the neural networks were used to classify the normality of reactions for a new dataset of ten repetitions of each single knockout (3411 simulations in total). From this, we generated a ‘flux profile’ for each simulation; a binary string of either 0 or 1 for each reaction within that simulation, where 0 means normal behaviour, and 1 means abnormal. We would expect some trends through the dataset that correspond to different classes of phenotype observed after gene knockouts, given that metabolism is central to cells. Reactions for neural networks with less than 70% accuracy were removed, leaving 267 reactions with a mean accuracy of 93.6%. We applied PCA to reduce the flux profiles to two dimensions while retaining most of the variance, and visualised, as shown in Figure 3. Each point is the flux profile of a simulation, and the principle components correspond to the reduced dimensions of the reaction flux profiles.

**Figure 3:**
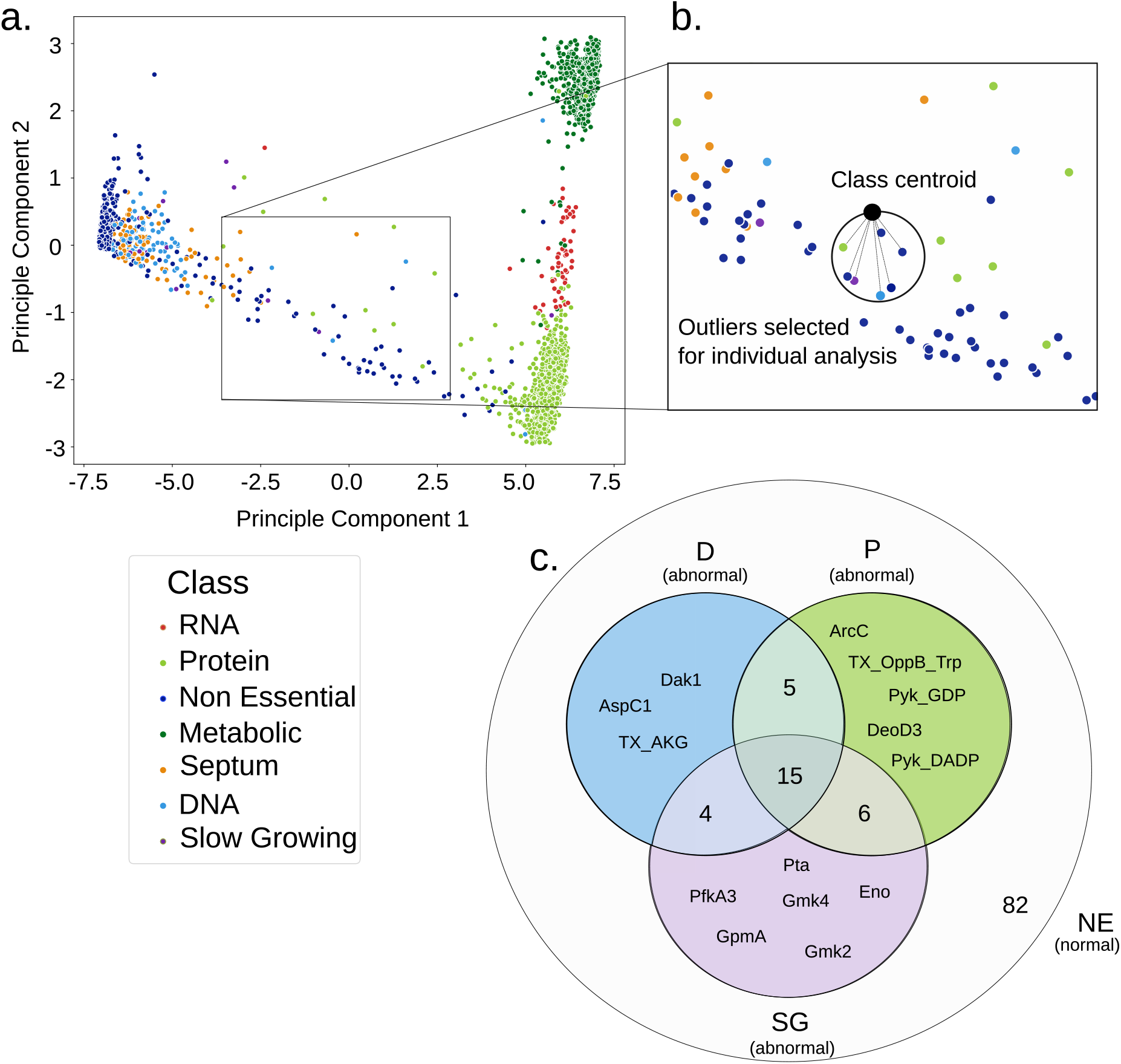
a) PCA plot of the flux profiles (binary strings of normal vs abnormal classifications for each flux in a simulation) from 3411 gene knockout simulations, reduced to two dimensions, then shown in different colours that correspond to manual labels. The classes are either defined by the presumed root cause of lack of cell division, or if the cell divides, the simulation is classified as non-essential. b) Seven simulations examined in further detail in the text are indicated. A schematic of the method used to choose the seven outliers is shown, where the centre of all of the class centroids is shown with its closest neighbours in a zoomed section of the PCA plot. c) Venn diagram showing the relationships between the reaction behaviour of the individual simulations chosen for analysis, to give biological context to the phenotype of each simulation. The reactions that behaved normally in all of the four non essential simulations (labelled as NE in the diagram, where the cell still divides after the gene knockout) were compared to the reactions that behaved abnormally in the three disrupted simulations — for the simulations from the DNA, protein and slow growing classes (labelled as D, P and SG respectively), the abnormally behaving set of reactions that do not intersect across the three simulations are listed and highlighted in the diagram. As these are three different phenotypic classes, it is the differences between them (and the non essential simulations) that we are most interested in. The numbers of shared reactions between the simulations, as well as those that behaved normally in all seven of the simulations, are shown but not listed individually.

The analysis dataset simulations were previously hand-labelled by phenotype according to differences in cell behaviour of the simulation output. The labelling classes were non-essential, DNA disruption, RNA disruption, metabolic disruption, protein disruption or septum disruption (Karr et al. 2012) — see Methods for details. The non essential class is defined by whether the cell divides or not, in keeping with current definitions of gene essentiality (Zhang & Zhang 2008), and the other classes are defined by what is indicated by the output data to be the root cause of cell death.

In Figure 3, several clear clusters of flux profiles are visible. To validate the significance of these groupings, we coloured the flux profile points according to manual labels of the phenotype that has occurred after the knockout; it can be seen (Figure 3) that these clusters correspond overall to the manually defined phenotypic classes. This suggests (intuitively) that it is different sets of reactions that behave abnormally for each different classes of phenotype, and/or significantly different scales in the amount of reactions affected, which will separate the different classes in the PCA space. We would expect the majority of reactions in a simulation that are labelled as non essential to be classified as normal, and the non essential simulations to be clustered together in the PCA space, as their flux profiles will be similar. Then, as we see simulations with greater disruption (e.g., the metabolic phenotypic class, where there is no growth, and no DNA, RNA or protein is created (see Methods section), where many reactions are behaving abnormally), these will be placed much further away from the non essential cluster.

### Individual simulation analysis of outliers

To understand the connection between the flux profiles and phenotypic classes of a simulation from a biological perspective, we chose several individual flux profiles to look at more closely. The entire dataset shows patterns in clustering of flux profiles that correspond with phenotypic classes, but the high dimensionality means we cannot infer specific meaning from how the metabolism relates to a phenotypic class without looking at individual reactions. Behaviour of certain reactions within a flux profile should give insight into why the phenotype for that simulation emerged, as we know that different reactions and metabolites are involved in different pathways, which are linked to different cellular processes.

In Figure 3, there are several outliers that diverge from their main clusters. To identify a set of simulations that capture some of the variance across the dataset, we found seven flux profiles that were closest to the mean of all of the class centroids (see Methods section* for more details), as the centre of the individual class centroids is a point in the PCA space that is far away from all of the clusters, and can be used to find outliers from multiple classes.

The chosen simulations (shown in Figure 3) contain four classified as non essential (where the cell divides), and one each of protein, DNA and slow growing phenotypes. Their flux profiles lie far away from others of their phenotypic class, so they clearly differ from the rest of their class, but still show the same output phenotype. Notably, the flux profiles are more similar to their non essential neighbours than to the simulations that they share their behaviour with, providing a basis to analyse the key differences between these simulations. For the three simulations with protein, DNA and slow growing phenotypes, we found the reactions that were abnormal in that simulation and normal in the other six, assuming that the surrounding four non essential simulations provide an approximate example of normal behaviour. The different sets of these reactions are shown in Figure 3.

In the flux profile of the DNA phenotype simulation, there are three reactions that behave abnormally whilst behaving normally in the other six simulations. These are AspC1 (conversion of L-Aspartate and 2-Oxoglutarate to L-Glutamate), Dak1 (phosphorylation of deoxyadenosine to create dAMP) and TX AKG (transport of 2-Oxoglutarate into the cell). Dak1 is the most relevant of the three to the end results — deoxyadenylate kinase catalyses an important step in the purine salvaging pathway (Griffith & Helleiner 1965) which contributes to DNA synthesis (Bizarro & Schuck 2007), so it may be the cause of the DNA phenotype. The other two reactions are both involved in metabolism of amino acids, and linked to each other — TX AKG is the transport of 2-Oxoglutarate, and then AspC1 is the conversion of L-Aspartate and 2-Oxoglutarate to L-Glutamate (Mavrides & Orr 1975). Although these are both steps in the nitrogen metabolism of *Mycoplasmas*, where the amino acids will later donate nitrogen to other compounds in the cell (Viljoen et al. 2013), in the model *M.genitalium* is cultured on rich media, where amino acids are available and transported into the cell, so their synthesis is not essential (Baseman et al. 2004). In this simulation, the reaction transporting glutamic acid into the cell via ABC transport is still behaving normally, indicating that this could be alleviating any problems caused by the malfunctioning of AspC1 and TX AKG.

In the protein phenotype simulation, the five reactions that behave abnormally are ArcC (carbamate kinase, which catalyses the synthesis of carbamoyl phosphate), DeoD3 (phosphorylation of deoxyguanosine), Pyk DADP, Pyk GDP (both conversions of phosphoenolpyruvate to pyruvate, generating dATP and dGTP respectively), and TX OppB Trp (transport of tryptophan into the cell). In this case, the tryptophan transport reaction may best explain the end phenotype — as *M.genitalium* does not synthesise tryptophan in its metabolism, if the transport reaction is disrupted, the cell will not be able to synthesise proteins as it will be missing an amino acid.

For the slow growing simulation, the abnormally behaving reactions are Eno (the conversion of 2-phosphoglycerate to phosphoenolpyruvate), Gmk2, Gmk4 (both phosphorylation of GMP), GpmA (internal transfer of phosphates), PfkA3 (phosphorylation of fructose) and Pta (phosphoryation of Acetyl-CoA). These reactions do not directly synthesise anything that is incorporated into other parts of the cell, which may explain why the cell still begins division even though its metabolic growth is impeded.

Although Figure 3 shows the variation in flux profiles across the dataset, there will be some circumstances where analysis of a single simulation is necessary. By comparing a flux profile to its nearest neighbours in the PCA space, we can see the main differences between simulations, and isolate reactions whose behaviour may be particularly relevant to the phenotypic class. For each of the chosen simulations described above, we have explored individual reaction behaviours within them, without having to consider the entire set of reactions, and seen that there is some connection between them and the phenotypic class that their simulations show.

### Analysis and biological context within the metabolic network

To ascribe biological meaning to trends seen across the dataset, we analysed the topology of the metabolic network, and how it is affected by reactions behaving abnormally after knockouts. The topology of metabolic networks has previously been found to be informative, where it was shown that the modularity of the *E.coli* metabolic network corresponds to metabolic functions (Ravasz et al. 2002), and so from a graphical perspective, we aimed to explain some of the biology behind the phenotypic classes and the flux profiles. The *M.genitalium* metabolic network is significantly smaller than many bacterial metabolisms (645 reactions vs e.g. 2382 in *E.coli* (Feist et al. 2007)), due to its genome size — however, analysis is not trivial. We used a graphical representation of the network, where each reaction is a node, and substrates that connect reactions are edges, as in the stoichiometric matrix of the metabolism in the knowledge base of the *M.genitalium* model. We visualised the reactions affected across each class in individual graphs, shown in Figure S2.

From the graphical representation, we used a maximal matching algorithm to find driver nodes. Driver nodes are the set of nodes in the metabolic network that need to be managed in order to have full control over the system (Liu et al. 2011) — therefore, in terms of input into the metabolism and flow through the metabolic pathways, their behaviour affects other reactions downstream, and they could be indicators of what might go wrong after gene knockouts. The driver nodes of the network are shown and named in Figure S3, with the whole metabolic network in graph form.

For each driver node, the pathways associated with that reaction were found from KEGG (Kanehisa et al. 2002), or (if there was no annotation for that reaction) the pathways associated with reactions that were one degree away from the driver, as shown in Table 3. If driver nodes are an indicator of the most important points of transport into the metabolism, the pathways that flow from them can show the cellular processes that would be disrupted if their input signals differ from wild-type behaviour.

**Table 2:** Manual labels of phenotypic classes (shown on the left-hand column) and their corresponding combinations of substance production (the column headings). A cross means that there is no active production of that substance in the case of DNA, RNA, and protein, and in the case of growth and division, these things do not occur. A tick means that opposite — so for example, in a simulation classified as ‘Non Essential’, we see production of DNA, RNA and protein, as well as both growth and division, and in a simulation classified as ‘Metabolic’, we see none of these things.In the case of the Slow Growing phentype, division begins at the end of the simulation but does not complete.

|  | DNA | RNA | Protein | Growth | Division |
| --- | --- | --- | --- | --- | --- |
| Metabolic | × | × | × | × | × |
| RNA | ✓ | × | × | × | × |
| Protein | ✓ | ✓ | × | × | × |
| Slow Growing | ✓ | ✓ | ✓ | ✓ | × |
| DNA | × | ✓ | ✓ | ✓ | × |
| Septum | ✓ | ✓ | ✓ | ✓ | × |
| Non Essential | ✓ | ✓ | ✓ | ✓ | ✓ |

**Table 3:** List of all of the driver nodes, whether they can linearly separate different phenotypic classes in the PCA space, and their associated pathways (if available).

| Driver | Pathways | Linearly separable |
| --- | --- | --- |
| TX_CO2 | Glycolysis, TCA cycle, Pyruvate metabolism, Carbon metabolism | Yes |
| TX_COA | Glycolysis, TCA cycle, Pyruvate metabolism, Carbon metabolism, Pantothenate and CoA biosynthesis, Methane metabolism | Yes |
| TXPYDX | Vitamin B6 pathway | Yes |
| TX_ACAL | Pentose phosphate pathway | No |
| TX_CAP | Purine metabolism, Carbon metabolism | No |
| TX_DDCA | Glycerolipid metabolism | n/a |
| TX_FOR | One carbon pool by folate, Carbon metabolism | n/a |
| TX_H2O2 | n/a | Yes |
| TX_HDCA | Glycerolipid metabolism | Yes |
| TX_HDCEA | Glycerolipid metabolism | Yes |
| TX_LIPOATE | n/a | Yes |
| TX_NAC | Nicotinate and nicotinamide metabolism | Yes |
| TX_O2 | Purine metabolism, Pyrimidine metabolism | Yes |
| TX_OA | Pyruvate metabolism, Carbon metabolism, Methane metabolism | No |
| TX_OCDCA | Glycerolipid metabolism | Yes |
| TX_OCDCEA | Glycerolipid metabolism | Yes |
| TX_RIBFLV | Riboflavin metabolism, Biosynthesis of secondary metabolites | Yes |
| TX_THF | One carbon pool by folate, Folate biosynthesis | Yes |
| TX_TTDCA | Glycerolipid metabolism | n/a |
| TX_TTDCEA | Glycerolipid metabolism | n/a |
| Upp | Pyrimidine metabolism | Yes |

Metabolic networks are known to be robust (Smart et al. 2008, Holme 2011), so many reactions can be individually removed without causing adverse effects. However, within *M.genitalium* metabolism, very few metabolites are organically synthesised (Dybvig & Voelker 1996). Transport reactions for essential substrates such as amino acids are far more important than they might be in a larger cell that has the capabilities to synthesise these things itself.

We found several driver nodes that can be individually used as features to divide the data into separate classes. For all driver nodes, we modelled a linear support vector machine (SVM) across the 2D data of the analysis dataset. We then selected those that could separate the data into normal vs abnormal behaviour with over 95% accuracy as good and simple indicators of metabolic behaviour, shown in Figure 4. 83% of the driver nodes could linearly separate the data with greater than 95% accuracy (listed in Table S3), compared with only 60% of the non-driver nodes, demonstrating their significance. Additionally, we can use individual driver nodes to mark phenotypic classes — normal behaviour for TX NAC, the reaction that transports nicotinamide into the cell, correlates strongly with the simulations classified as non-essential, with a phi coefficient (a measure of correlation between binary variables (Ekström 2011)) of 92%. Behaviour of TX RIBFLV can split the dataset into the classes where we see growth (Non essential, Septum and DNA phenotypes) and the classes where there is no growth (Metabolic, RNA and Protein phenotypes) with a phi coefficient of 95%. Equally, we can see that abnormal behaviour for Upp (dephosphorylation of uracil) is strongly indicative of a metabolic phenotype, and can be used as a feature to separate metabolic disruption phenotypes from other types of phenotype, with a phi coefficient of 97%. Overall, the driver node analysis showed that it is possible to identify important reactions within the network that correlate with certain cell behaviour, meaning that we can focus on these to understand the end phenotype rather than the entire set of reactions.

**Figure 4:**
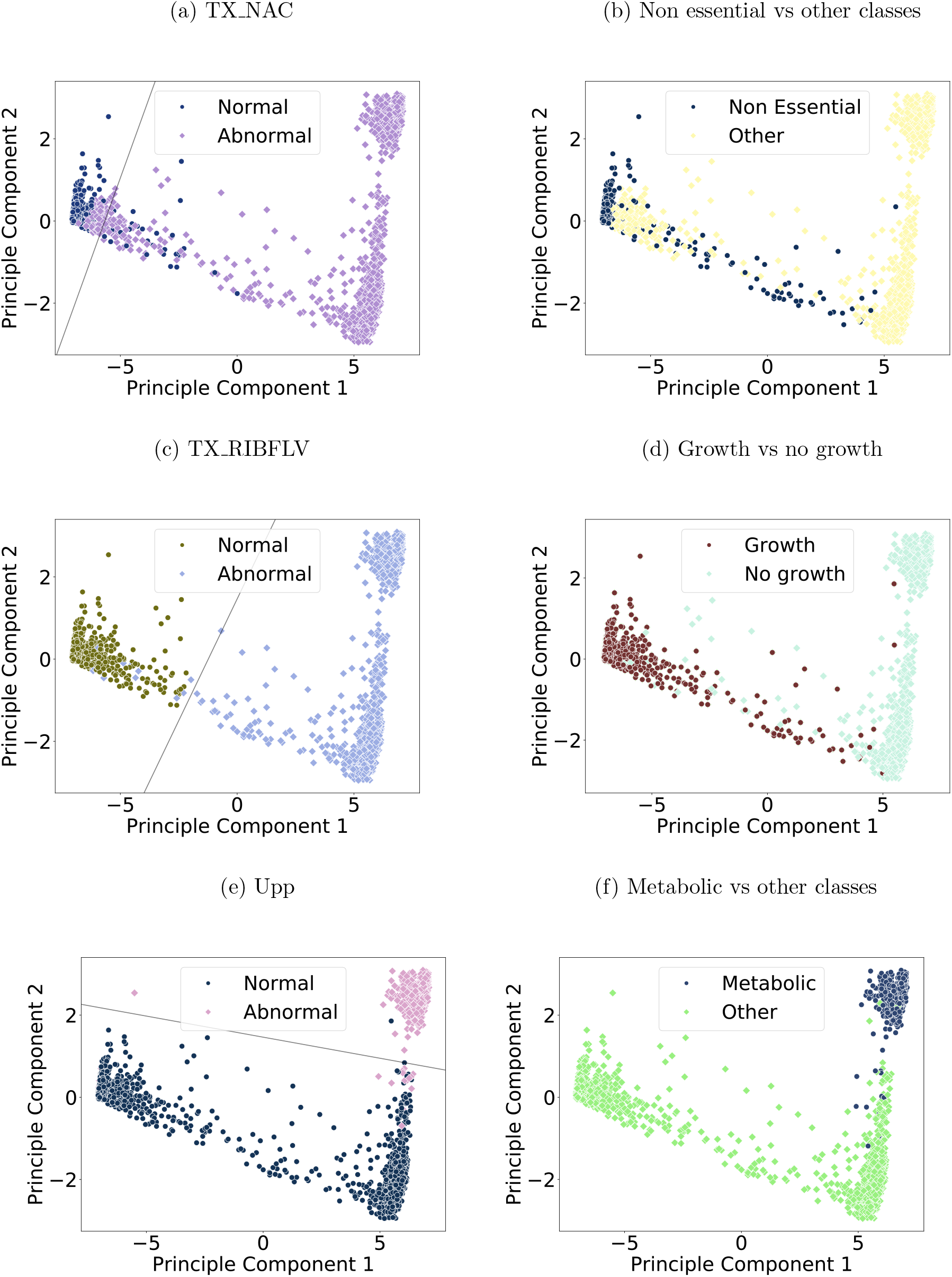
Scatter plots of all flux profiles reduced to a 2D feature space. On the left (plots a,c, and e), each point is labelled with the behaviour (normal or abnormal) of a single reaction that is a driver node. The lines shown decision boundaries for support vector machines (SVMs); models that form a hyperplane to linearly separate different classes of data, where in this case the classes will be flux profiles where the specified reaction behaves normally, and flux profiles where the specified reaction behaves abnormally. On the right (plots b, d and f), each point shows the manual label of the phenotypic classes that correlate with the behaviour of the reactions on the left — the simulations that show normal behaviour of TX NAC have a 92% correlation with those that are manually classified as non essential; those that show normal behaviour of TX RIBFLV show a 95% correlation with those that are manually classified as Non essential, DNA or Septum phenotypes (which all show cell growth) and those that show abnormal behaviour for Upp show a 97% correlation with those that are manually classified as Metabolic.

## Discussion

The processing of complex data is imperative to understanding whole-cell model output. We have shown a variety of different analysis methods that can take a large high-dimensional dataset and distill it into visualisations that are easy to interpret, showing how the behaviour of specific reactions can be used as marker of a particular phenotypic class and their importance to the corresponding cellular process.

Understanding the effects of single gene knockouts is a deceptively difficult task, as the domino effect of gene removal can cause large changes in the behaviour of a cell through its life cycle. Visualising and analysing thousands of time series is a challenge faced by many branches of research. These two problems come together in the context of whole-cell models. Using Snorkel and neural networks, we have been able to classify metabolic fluxes as normal or abnormal and visualise them in two dimensions, meaning that the dataset separates out into groups that can be interpreted. Whole-cell model data must be understood in the context of controllable biological mechanisms to be relevant to genome design: in order to use knowledge gained from modelling in real cells, we must understand the internal operations as well as the output. The flux behaviour across different gene knockouts, and in particular the driver nodes, can show the links between genotype and phenotype, plus unprecedented effects that a gene may have on reactions seemingly unrelated to its functional annotation, on a scale that is only possible in a whole-cell model. As this analysis gives an overview of the entire metabolism, we can approach the problem of understanding gene knockouts in a way that includes the genomic context of the remaining genes and the behaviour of their associated reactions, rather than examining the phenotype with regards to the single gene that has been removed.

For genome design, there has long been an idea of ‘modularity’ in cells, at different scales and abstractions (Papin et al. 2004). Cellular subsystems that use a unique set of molecules and rules to perform a function such as DNA replication or glycolysis use chemical specificity to keep their processes separate from other functional modules (Hartwell et al. 1999). It has been proposed recently that the future of genome design may be in minimal cells, combined with different functional modules to create cells for specific purposes (Gibson 2014). This would require a detailed understanding of not only how a genome maps to its phenotype and how the genes themselves can form functional modules, but also concerning the ways in which these modules interact.

It has been hypothesised by Ritchie *et al.* that the connections between different functional modules could either be linear (where changes in the genome filter down through the transcriptome and proteome to the phenotype) or nonlinear (where all functional modules can contribute in individual ways to the end phenotype (Ritchie et al. 2015)). In the *M.genitalium* model, the 28 submodels are all connected and integrated together at each timestep, by calculating the amounts of different molecules formed and allocating them to the submodels according to equations and parameters for rates of production and degradation. The model is both nonlinear and stochastic, where small variations in concentrations of substances can elicit large changes for other processes during the cell life cycle. In this sense, it may be a more exploratory proxy for cell behaviour than CBMs and their more complex successors (O’brien et al. 2013, Du et al. 2019), as these use systems of linearised differential equations to make deterministic predictions.

The metabolism in the *M.genitalium* model is linked to all of the other submodels, and is a central hub of activity. Regardless of any hierarchical structure, the metabolism is an integral stepping stone for substance transfer between cellular processes. Although internal mechanisms and local rules for the model were gathered from experimental data and are biologically valid, the complexity that arises from so many parameters being integrated together means that the model has to be treated as a black box. Analysing the behaviour of the model could ultimately lead to better biological understanding of the connections between cellular processes. If the way that two processes are coupled together *in silico* in the whole-cell model yields output that matches experimental data, this can help to develop insight into how these processes are linked in a real cell. This could aid genome design, where insights from modelling can rationally guide *in vitro* experiments and gene editing (Landon et al. 2019, Rees-Garbutt et al. 2020).

We can see from Figure 3 that the gene knockouts that cause DNA and septum disruptions cause similar behaviour in the flux profiles to non-essential gene knockouts, as their clusters are very close together, which is likely because the majority of their reaction behaviour was classified as normal. Additionally, Figure S2 shows us that fewer than ten reactions are consistently affected across the simulations within these phenotypic classes, and so we can infer that these reactions are probably the bridge between the metabolism process and the DNA replication or cytokinesis process. Limitations of the *M.genitalium* model mean that the results presented here do not include multiple cell divisions and it is possible that more widespread effects on metabolism would be revealed in future work with more cell generations.

The interactions between the metabolism and the other phenotypic classes (protein and RNA) are less simple, as there are significantly more reactions that are consistently behaving abnormally in these simulations. This is not surprising, as there are two main functions for a cell to perform: growth and replication. Growth occurs consistently through the cell cycle, and requires constant synthesis and degradation of different proteins and RNAs. There is also a temporal element, as cascades of reactions that form different proteins may need to occur in a specific order. Any disruption to an aspect of this process during the life cycle will filter down to other processes, whereas if DNA replication is disrupted, it is primarily cell division that will be halted. In future studies, it would be interesting to see if dividing the proteins into functional groups and pathways for further analysis leads to better understand their roles and how they interact with each other.

There are limitations to this type of analysis: it is hard to draw solid conclusions about cell behaviour when *M.genitalium* is an organism where not all of the genes are classified, and the data that the model was built upon is from many different sources and organisms. In terms of the network analysis, there are some reactions that, although they have been observed in *M.genitalium*, do not have known enzymes to catalyse them, which leaves gaps within the model. There may be unexpected and unusual behaviours that are not captured in the training data as well, leading to misclassifications; for example, the reactions that performed badly in the neural network classifications may be sensitive to small changes in the metabolic network, meaning that their behaviour is inconsistent and unpredictable. However, it is useful to flag these reactions, and in the future use different approaches to understand their behaviour. There is also the possibility that, after applying the machine learning processes, the results show more about the internal features of the model itself than the actual biology, which is a good starting point for lab work.

As whole-cell models become more commonplace and widely used, analysis software will become more and more important. Understanding is likely to increase as more familiar organisms are modelled. The most recent whole-cell model is of *E.coli*, which is a better understood organism than *M.genitalium*, with significantly more data available to validate and add to it, so this is an important development for the field. However, the complexity of models will increase hugely with the size of the genome of the organism, and as *E.coli* has an order of magnitude more genes than *M.genitalium* (Blattner et al. 1997), analysis tools that can provide data processing and dimensionality reduction will be even more important for enhancing understanding, and ultimately genome design.

## Acknowledgements

We would like to thank the Advanced Computing Research Centre (ACRC) and BrisSynBio, a BBSRC/EPSRC Synthetic Biology Research Centre, at the University of Bristol for access to the BlueCrystal and Bluegem supercomputers. Special thanks to the HPC and RDSF teams of the ACRC, particularly Dr. Christopher Woods, Simon Burbidge, Matt Williams, and Damian Steer for their help with BlueCrystal, BlueGem, data storage and publication. We would like to thank Dr Thomas Gorochowski (University of Bristol) for useful feedback on the manuscript. We would like to thank Dr Joshua Rees-Garbutt for generating the single-gene KO simulations we analysed. L.M. is supported by the Medical Research Council grant MR/N021444/1, by the Engineering and Physical Sciences Research Council (grants EP/R041695/1 and EP/S01876X/1) and by the EU Horizon 2020 research project COSY-BIO (grant 766840); O.C., L.M. and C.G. are supported by a BrisSynBio, a BBSRC/EPSRC Synthetic Biology Research Centre (BB/L01386X/1), flexi-fund grant; O.C. is supported by the Bristol Centre for Complexity Sciences (BCCS) Centre for Doctoral Training (CDT) EP/I013717/1; S.L. is supported by EPSRC Future Opportunity Scholarships.

## Author Contributions

C.G., L.M., O.C., G.B. and S.L. were involved in the ideation. G.B. created the generative model and implemented Snorkel, with help from O.C. S.L. trained the neural networks, carried out the network analysis, created the figures and wrote the paper. C.G., L.M., O.C. and J.R. were involved in editing and feedback on paper.

## Competing interests

The authors declare no competing interests.

## STAR Methods

### Description of the data

We began with two sets of data — one to train the machine learning models, and one to apply them and analyse the output. The simulations were generated from running the *M.genitalium* whole-cell model on a supercomputer cluster, with each gene singly knocked out. The model requires 8GB of RAM for each simulation, and was run on BlueGem, a 900 core supercomputer at the University of Bristol, using MATLAB R2013b. It is available at https://github.com/CovertLab/WholeCell. The raw metabolic flux timeseries was then converted to Pandas dataframes and stored in a .pickle format to save space. The training set consisted of time series of reaction fluxes for three repetitions of every possible single knockout from the *M.genitalium* model, of which there are 359, plus 200 wild-type simulations. Each time series is 50,000 seconds in total, and we used the time series of 279 reactions from each simulation. This was 1270 simulations, in total. The dataset that we applied the analysis to consisted of ten repetitions of all of the single gene knockouts, with the same reaction time series, and so this dataset was 3411 simulations in total. One knockout, MG 469, consistently caused the model to crash and simulations to terminate, and a few simulations did not complete due to errors on the supercomputer cluster.

### Labelling

Snorkel is a system that takes input data points and manually defined labelling functions, and collates these into a generative model that outputs probabilistic labels for the data. The labelling functions will produce noisy labels, which are then used as weak supervision for a stronger predictive function by combining three measures — the labelling propensity (whether the data point has been assigned a label or not), the accuracy of each label, and the correlations of the multiple labelling functions. The label matrix generated from these measures is then used to define an exponential distribution which can predict probabilistic training labels. The normality of ten reactions was manually labelled by visual inspection of the time series, comparing features of the plots such as smoothness and linearity to wildtype time series from the same reactions (Correll & Heer 2017, Correll et al. 2012), and used to validate Snorkel’s weak labels; the accuracies of which are shown in Table 1. The algorithm was implemented using the snorkel library in Python.

The manual labelling was done based on the phenotypic classes defined by the original publication of the *M.genitalium* model, which used the production capacity of various features from the model output to classify a simulation (Karr et al. 2012). The combinations of these features that contribute to a particular class are detailed in Table 2, and the simulations used in the analysis dataset were all labelled by manual inspection of the model output.

### Training and tuning the neural networks

Once the data is fully labelled, a standard discriminative model can be trained for classification. In this case, we chose to use a neural network, implemented with the Python library tensorflow (version 2.0.0-rc0). Using the data labelled by the generative model, a neural network was trained for each reaction. Different combinations of hyperparameters (epoch size, batch size, and number of nodes in a layer) were tested, so that an optimal combination could be used for each network to find the highest accuracy. Generally, the combination of hyperparameters can have a significant effect on the neural network output, so these factors are important. Epoch size refers to the number of rounds of back-propagation performed by the network, batch size means the number of training data samples input before the model updates, and number of nodes refers to number of nodes of the network in each hidden layer. Epoch size will leave the data underfitted if too small, and overfitted if too large; batch size is generally optimised for processing time (in that larger batch sizes will train the network faster, whereas a smaller batch size may help the weights converge faster); and number of nodes is usually taken to be some number between the amount of input nodes and the amount of output nodes. There is no set method for selecting hyperparameters for neural nets, and it is frequently taken to be a trial-and-error process (Sarle 1994). We tuned our neural networks via a brute-force approach, where different parameters within a set range were trialled to increase the accuracy of the network. Epoch size was kept relatively low, as after some testing, many of the neural networks converged to accuracies > 95% after only five epochs, and so we tested epoch values of 5, 10 and 15. Batch sizes of 50, 100 and 150 used, and node numbers of 750, 1500 and 2250 were tried, where we selected the network with hyperparameters that gave the highest accuracy. The reactions from neural networks that gave accuracy of less than 70% were removed, leaving 267 reactions and neural networks with a mean accuracy of 0.936%.

### Selection of individual simulation outliers

To choose outlier simulations from the PCA space, we first found the centre of each of the seven different classes by taking the mean of all of the positions of the points within that class. We then found the centroid of these seven centres, leaving us with a single point that is far from all of the clusters. We went on to iteratively find the points with the smallest Euclidean distance from the centroid, stopping when four different classes of simulation had been selected.

### Network formation and features

After the neural networks were trained and fluxes classified across the dataset, we turned to network analysis. With the stoichiometric matrix for the metabolism, **S**, taken from the *M.genitalium* model knowledge base, we formed a metabolic graph **A** from the relationship

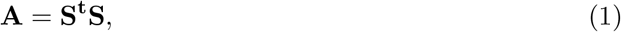

which creates a widely used graphical representation of a metabolic network, where the reactions form nodes of the graph, and the substrates form edges that connect them (Palsson 2006). We were able to find a set of driver nodes (the set of nodes that must be controlled in order to fully control the network) using the maximal matching function in Python’s networkx library (version 2.4). The pathways associated with the driver nodes were found via the EC numbers from the supplementary material of the *M.genitalium* model (Karr et al. 2012), where the Python library bioservices was used to look up the pathways for each EC number from KEGG.

The metabolic sub-networks were plotted in python-igraph where, for each class across the dataset, the affected reactions are shown as a sub-network with a colour gradient corresponding to how frequently that reaction behaves abnormally. A threshold for ‘noisy’ reactions was found from wild-type simulations, where an exponential distribution was fitted to the frequencies of reactions classified as behaving abnormally by the neural networks. For a wild-type simulation, in theory all reactions should be classified as normal, but, as the *M.genitalium* model is stochastic, there can be a range of different behaviours, depending on the initial conditions of the simulation and other random processes (e.g. radiation and DNA damage). The interval under which 95% of the data was contained was found, and this value was selected as a rate parameter, which was used as the threshold of significance for whether a reaction was considered to be behaving abnormally consistently.

We then performed PCA using the SciPy library (version 1.3.1) to reduce the data to two dimensions and plot the data on a scatter plot using Seaborn (version 0.9.0). After the reduction, 84% of the variance in the full dimensions of the data was conserved, so there was not significant information loss after this operation. After having found the driver nodes, we trained a linear support vector machine (SVM) for the normality of each one to separate the data points on the PCA plot, selecting those that could divide the data with *>* 95% accuracy. For the SVM, we used the sklearn Python library (version 0.21.3).

## Notes

### Competing Interest Statement

The authors have declared no competing interest.

