## Supplementary information for "Understanding metabolic behaviour in whole-cell model output"

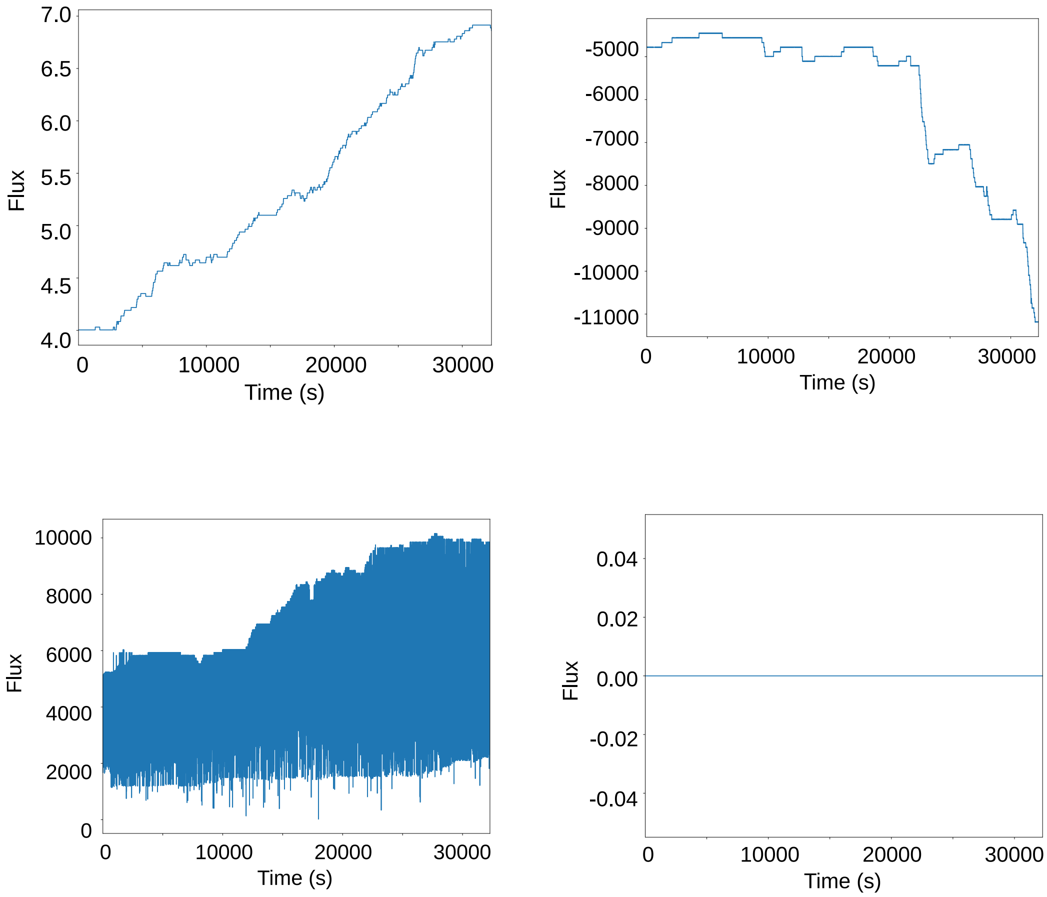


Figure S1: Examples of different metabolic flux behaviours (clockwise from top left: increasing (Pyk_GDP), decreasing (DcdK), oscillatory (AceE), stationary (TX_m1dG) for different reactions across the entire timeseries of a single simulation.

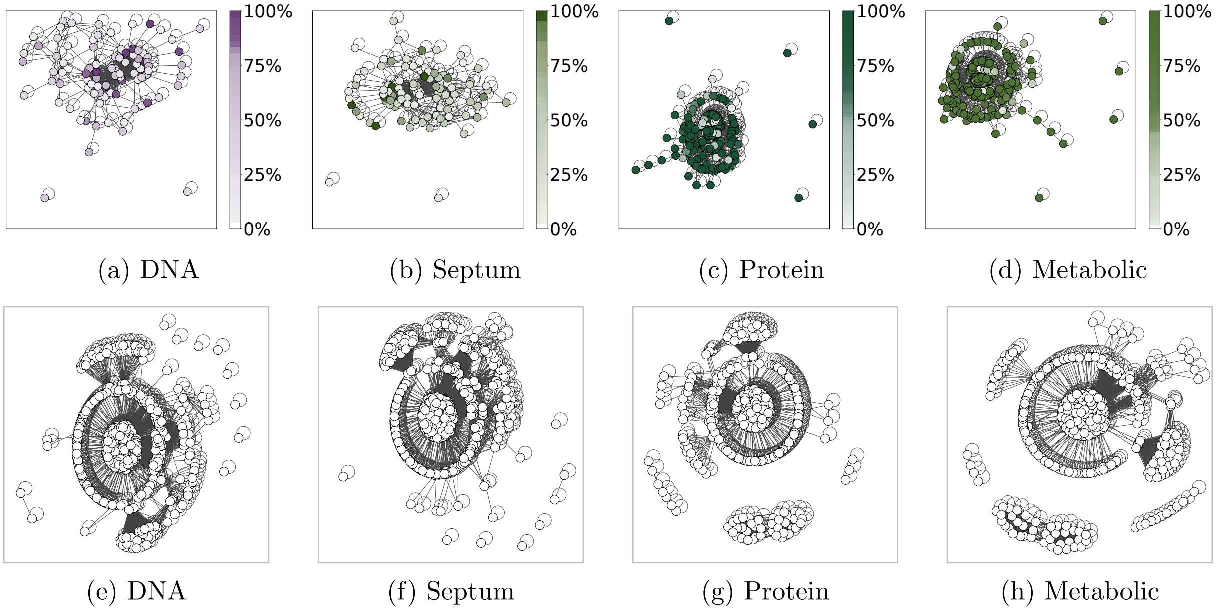


Figure S2: Sub-networks of the behaviour of reactions across the simulations within individual classes (shown here are the DNA, Septum, Protein and Metabolic phenotype classes). The reactions that are consistently behaving abnormally are shown on the top row, with a colour gradient that corresponds to how frequently the reaction behaves abnormally, and the remaining reactions (either those that behave normally, or are unclassified) are shown on the bottom row.

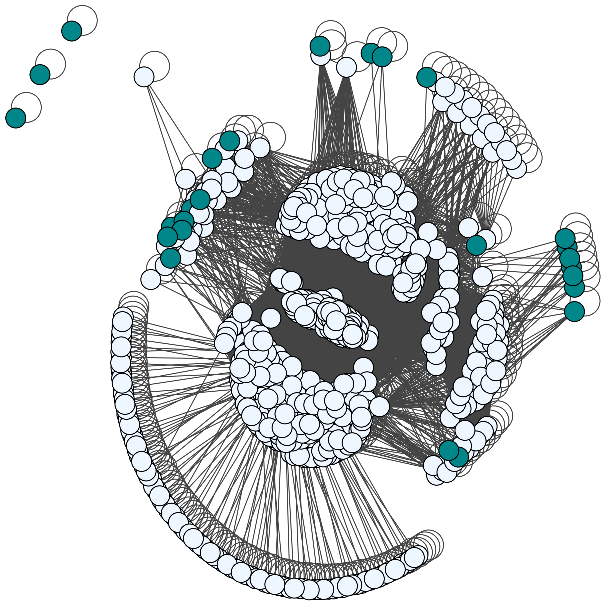

| Driver Node |
|---|
| TXPYDX (Pyroxidal ) |
| TX_ACAL (Acetaldehyde ) |
| TX_CAP (Carbamoyl phosphate ) |
| TX_CO2 (Carbon dioxide ) |
| TX_COA (Coenzyme A ) |
| TX_FOR (Formate ) |
| TX_H202 (Hydrogen peroxide ) |
| TX_HDCA (Hexadecanoate ) |
| TX_HDCEA (Hexadecenoate ) |
| TX_DDCA (Dodecanoate) |
| TX_LIPOATE (Lipoate) |
| TX_NAC (Nicotinade) |
| TX_O2 (Oxygen) |
| TX_OA (Oxaloacetate) |
| TX_OCDCA (Octadecanoate) |
| TX_OCDCEA (Octadecenoate) |
| TX_RIBFLV (Riboflavin) |
| TX_THF (Tetrahydrofolate) |
| TX_TTDCA (Tetradecanoate) |
| TX_TTDCEA (Tetradecenoate) |
| Upp (Uracil dephosphorylation) |

Figure S3: The metabolic network shown in graph form, where the nodes are reactions and edges are substrates that connect them (e.g. if a substrate is involved in two different reactions, this becomes an edge between the two reaction nodes). The driver nodes are shown in dark turquoise, which are the set of nodes that need to be controlled in order to have full control over the network. The driver nodes are listed in the adjacent table.
